# Ångström-resolution imaging of cell-surface glycans

**DOI:** 10.1101/2025.02.07.637003

**Authors:** Luciano A. Masullo, Karim Almahayni, Isabelle Pachmayr, Monique Honsa, Larissa Heinze, Sarah Fritsche, Heinrich Grabmayr, Ralf Jungmann, Leonhard Möckl

**Affiliations:** Max Planck Institute of Biochemistry, Planegg, Germany; Max Planck Institute for the Science of Light, Erlangen, Germany; Department of Physics, Friedrich-Alexander-University Erlangen-Nuremberg, Erlangen, Germany; Faculty of Physics and Center for Nanoscience, Ludwig Maximilian University, Munich, Germany; Faculty of Medicine 1/CITABLE, Friedrich-Alexander-University Erlangen-Nuremberg, Erlangen, Germany; Deutsches Zentrum Immuntherapie, University Clinic Erlangen

**Author notes:** These authors contributed equally (Luciano A. Masullo, Karim Almahayni).

## Abstract

Glycobiology is rooted in the study of monosaccharides, Ångström-sized molecules that are the building blocks of intricate glycosylation patterns. Glycosylated biomolecules form the glycocalyx, a dense coat encasing every human cell with central relevance – among others – in immunology, oncology, and virology. In order to understand glycosylation function, visualizing its molecular structure is fundamental. However, the ability to visualize the molecular architecture of the glycocalyx has remained elusive. Techniques like mass spectrometry, electron microscopy, and fluorescence microscopy lack the necessary cellular context, specificity, and resolution. Here, we address these limitations by combining metabolic labeling with Ångström-resolution fluorescence microscopy, enabling the first-ever visualization of individual sugars within glycans on the cell surface. Our work provides unprecedented insights into the molecular architecture of the glycocalyx and constitutes the foundation for future explorations of its function in health and disease.

## Introduction

The glycocalyx covers all cells in the human body. It is composed of glycosylated proteins, glycolipids, free polysaccharides, and the recently discovered glycoRNAs (*1, 2*). The glycocalyx plays a fundamental role in a range of cellular processes in health and disease, including immune system regulation (*3*), cell signaling (*4, 5*), leukocyte adhesion (*6*), and cancer development (*7*).

A critical challenge in glycobiology has been the structural analysis of cell-surface glycans, which are the fundamental components of the glycocalyx at the molecular level. This question has been addressed by a number of advanced characterization methods, in particular, mass spectrometry (*8*), scanning tunneling microscopy (*9*), electron microscopy (*10*), and light microscopy (*11*). Each method revealed important aspects of cell-surface glycosylation. Mass spectrometry-based methods were used to infer glycan structures (*12*); scanning tunneling microscopy allowed for the visualization of isolated free glycans and glycans attached to proteins and lipids (*13*), and light as well as electron microscopy enabled measurements of glycocalyx thickness in tissue sections and cultured cells (*11, 14*).

However, each of these methods has limitations. Mass spectrometry requires the removal of glycans from the cell surface for ionization(*8*); scanning tunneling microscopy also requires the isolation of glycans (*13*); electron microscopy sample preparation is damaging to the glycocalyx and lacks species-specific contrast (*15, 16*). Fluorescence microscopy would be an ideal method to study the glycocalyx *in situ* due to its low invasiveness and cellular compatibility. In order to spatially analyze glycocalyx structure with molecular specificity in cells using fluorescence microscopy, one key ingredient is efficient and specific tagging of target sugars. However, the field of glycobiology has traditionally suffered from the limited availability of methods to specifically and efficiently label structural building units, compared to DNA, RNA or protein biology. In particular, there is a limited availability of antibodies against glycan structures, which, in addition, often exhibit low affinity (*17*). Furthermore, genetic labeling approaches are not applicable, as glycans are secondary gene products and therefore not directly encoded in the genome (*18*). Finally, lectin-based glycan labeling can be used, however, lectins show rather poor affinity and specificity (*19*).

Recently, the discovery of metabolic incorporation of unnatural sugar analogs in conjunction with live-cell click chemistry has provided the field of functional glycobiology with an asset to specifically label individual sugar residues within the glycocalyx. This ability has been used for fluorescence microscopy (*20, 21*) and even super-resolution approaches (*11, 22*). However, the expected distances between individual sugars (often densely packed within the glycocalyx) based on available structural studies (*23, 24*) are at the sub-10 nm scale and even below one nanometer. While in principle click-chemistry holds the potential to achieve sub-nm resolution due to the size of the labeling molecule (8 Å), conventional super-resolution microscopy methods like (d)STORM (*25, 26*), lack the spatial resolution to resolve the molecular architecture of the glycocalyx (*27*). While previous studies enabled relevant insights (*11, 28*), they still essentially lacked the resolution to resolve details below 20 nm. Moreover, recent reports (*29*) indicate that, at distances below 10 nm, photophysical interactions between fluorophores significantly influence fluorescence emission, thereby limiting super-resolution techniques that rely on labeling the imaging target with a single, fixed dye to resolutions of approx. 10-20 nm (*30*). A method to study cell-surface glycans at molecular resolution within the native cellular context is missing so far.

We have recently introduced RESI (Resolution Enhancement by Sequential Imaging) (*31*), a DNA-barcoding optical microscopy method that achieves Ångström-resolution. While Ångström resolution was demonstrated in DNA origami nanostructures, the achievable resolution in cells was experimentally limited to the size of the labeling probes, i.e. nanobodies of around 5 nm in size. Thus, achieving Ångström-resolution in cells has remained elusive and limited by the labeling approach.

Here, we combine RESI (*31*) with bioorthogonal metabolic labeling (*32, 33*), which allows us – for the first time – to resolve glycans down to individual sugars with Ångström resolution in whole cells. Leveraging this unique spatial information, we show that sugar residues form distinct spatial arrangements on the surface of cells that are smaller than the size of single proteins. In-depth quantitative analysis reveals previously elusive molecular signatures of single sugars within glycans on individual proteins. Taken together, we establish RESI combined with metabolic labeling as a transformative technique in glycobiology with the prospect of linking glycan structure to function, identifying molecular glycocalyx changes related to disease progression, discovering novel therapeutic avenues, and developing diagnostic tools. From a methodological perspective, our work constitutes the first demonstration of optical Ångström resolution in a native cellular context.

### RESI in combination with metabolic labeling enables Ångström-resolution imaging of cell-surface glycosylation

Monosaccharides, just a few Ångströms in size, serve as the building blocks of complex glycans and glycosylated biomolecules (**Fig. 1A**). To resolve these cell-surface carbohydrates, we combine efficient and specific metabolic labeling with the Ångström spatial resolution of RESI. First, we developed and optimized a method to attach single strands of DNA to the sugars of interest (**Fig. 1B**). RESI is based on DNA-PAINT (DNA Points Accumulation for Imaging in Nanoscale Topography) (*34, 35*) a super-resolution microscopy method that relies on the transient binding of fluorescently labeled DNA probes to complementary target sequences, achieving approximately 5-10 nm spatial resolution through single-molecule localization. By stochastically isolating and sequentially imaging sparse subsets of targets at this resolution (**Fig. 1C**), RESI increases the precision of DNA-PAINT measurements by averaging localizations (**Fig. 1D**), enabling spatial resolution at the Ångström level (**Fig. 1E**). This is critical to resolve individual glycans and individual sugars within glycans (**Fig. 1F**).

**Fig. 1.**
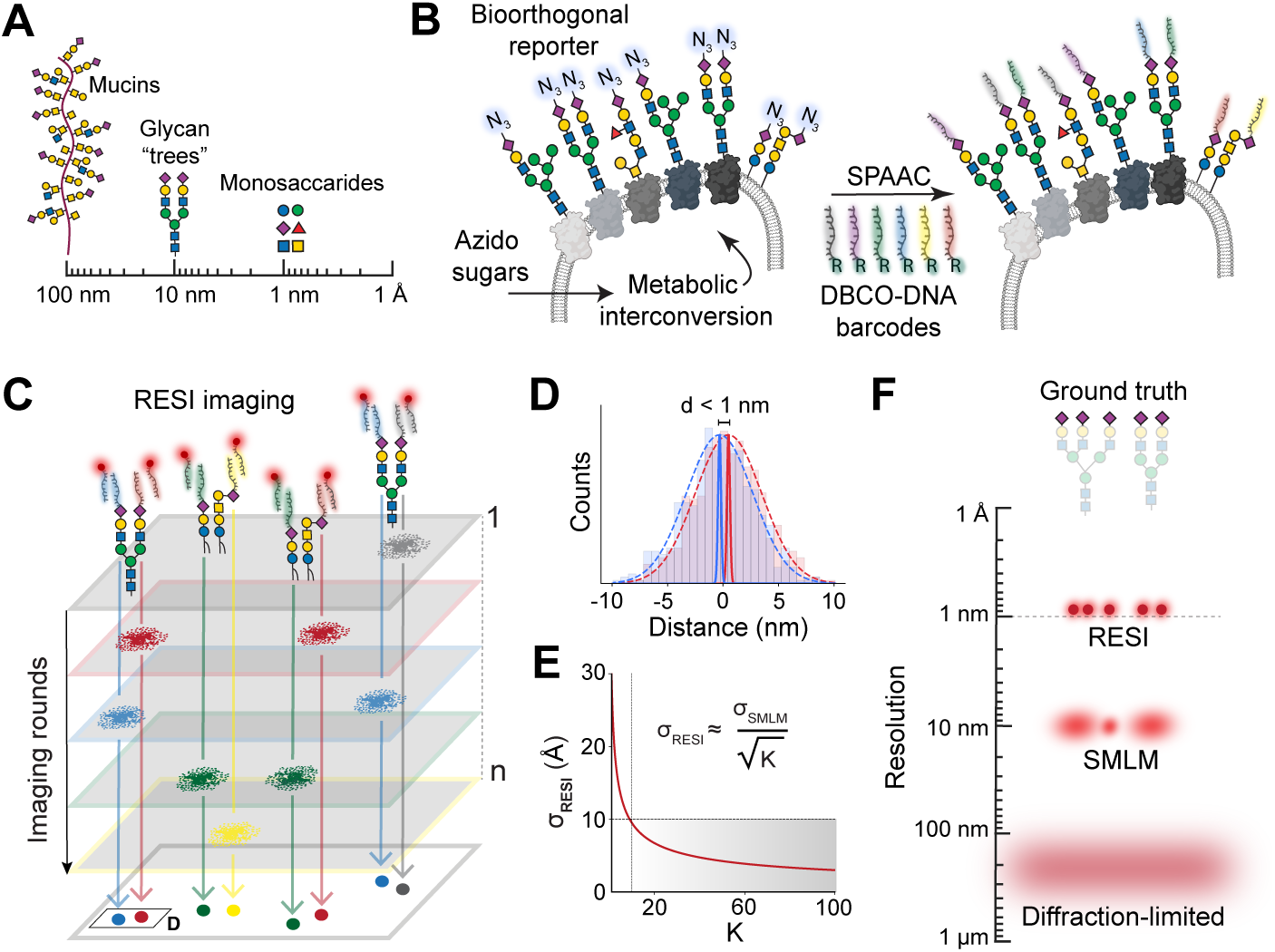
Experimental concept. (**A**) Monosaccharides represent the smallest length scale in glycobiology, forming the fundamental building blocks for larger glycan structures (tens of nanometers) and heavily glycosylated mucins (up to hundreds of nanometers). Monosaccharides are depicted following the Symbol Nomenclature for Glycans (SNFG) guidelines (*55*). (**B**) Azido sugars are metabolized by cells and integrated into target monosaccharides introducing a bioorthogonal azido group as a molecular reporter. This azido group facilitates the attachment of six orthogonal DBCO-modified DNA strands (shown in different colors) via a strain-promoted azide-alkyne click chemistry reaction (SPAAC), enabling precise labeling of the target monosaccharide unit. (**C**) Imaging sugar molecules, labeled with distinct DNA barcodes, through the sequential addition of their corresponding imaging DNA sequences, facilitates temporal separation of signals distinguishing blinks from nearby molecules. (**D**) Combining all localizations per target (𝐾) from each imaging round enhances localization precision. (**E**) In RESI, localization precision improves with 1⁄√𝐾, thus resolution enhancement is independent of 𝜎*_SMLM_*, achieving Ångström-scale precision. (**F**) Unlike other super-resolution imaging techniques, RESI is capable of resolving single sugars within a glycan. The glycan structures exemplify sialic acid labeling.

To enable RESI imaging of individual sugar residues within glycans, we optimized metabolic labeling to facilitate efficient attachment of DNA probes. In particular, we employed tetraacetylated N-acetylgalactosamine (Ac_4_GalNAz) to label N-acetyllactosamine (LacNAc; i.e., Galβ1-4GlcNAc) residues and tetraacetylated N-acetylmannosamine (Ac_4_ManNAz) to label N-acetylneuraminic acid (Neu5Ac) with azido sugars. In the following, Neu5Ac is referred to as “sialic acids” for simplicity. The azido sugars are subsequently covalently linked to six orthogonal dibenzocyclooctyne (DBCO)-modified RESI-compatible DNA strands via strain-promoted live-cell copper-free click chemistry (*33*) (**Fig. S1**). We opted for copper-free click chemistry, as it has been shown to achieve higher labeling efficiencies compared to copper-catalyzed click chemistry (*27*). The concentration of docking strands was optimized to ensure complete saturation of available azido groups (**Fig. S2**).

In the context of RESI’s Ångström-resolution capabilities, metabolic labeling offers the critical advantage of a much smaller labeling footprint (below one nanometer) compared to antibody (10-15 nm) and even nanobody-based labeling (5-10 nm) as the DNA strands are attached directly to the sugar target without the need for a genetically encoded tag or exogenous affinity reagents.

### Visualization of individual sugars *in situ* with Ångström-resolution

To demonstrate the capability of our approach, we imaged Human Microvascular Endothelial Cells (HMECs) where sialic acids were tagged with azido groups via Ac_4_ManNAz incorporation in order to benchmark spatial resolution using TIRF, STORM, DNA-PAINT, and RESI. First, we used cells labeled with DBCO-Alexa Fluor 647 and imaged them using diffraction-limited Total Internal Reflection Fluorescence (TIRF) microscopy (*36*) at ∼250 nm resolution. The same sample was then imaged with (d)STORM (*25, 26*) achieving ∼10 nm localization precision (∼25 nm resolution) (**Fig. 2A-D**). Despite offering spatial resolution an order of magnitude better than the diffraction limit (**Fig. 2B**), STORM fails to resolve the details of the sugar distribution at the true molecular scale (**Fig. 2D**).

**Fig. 2.**
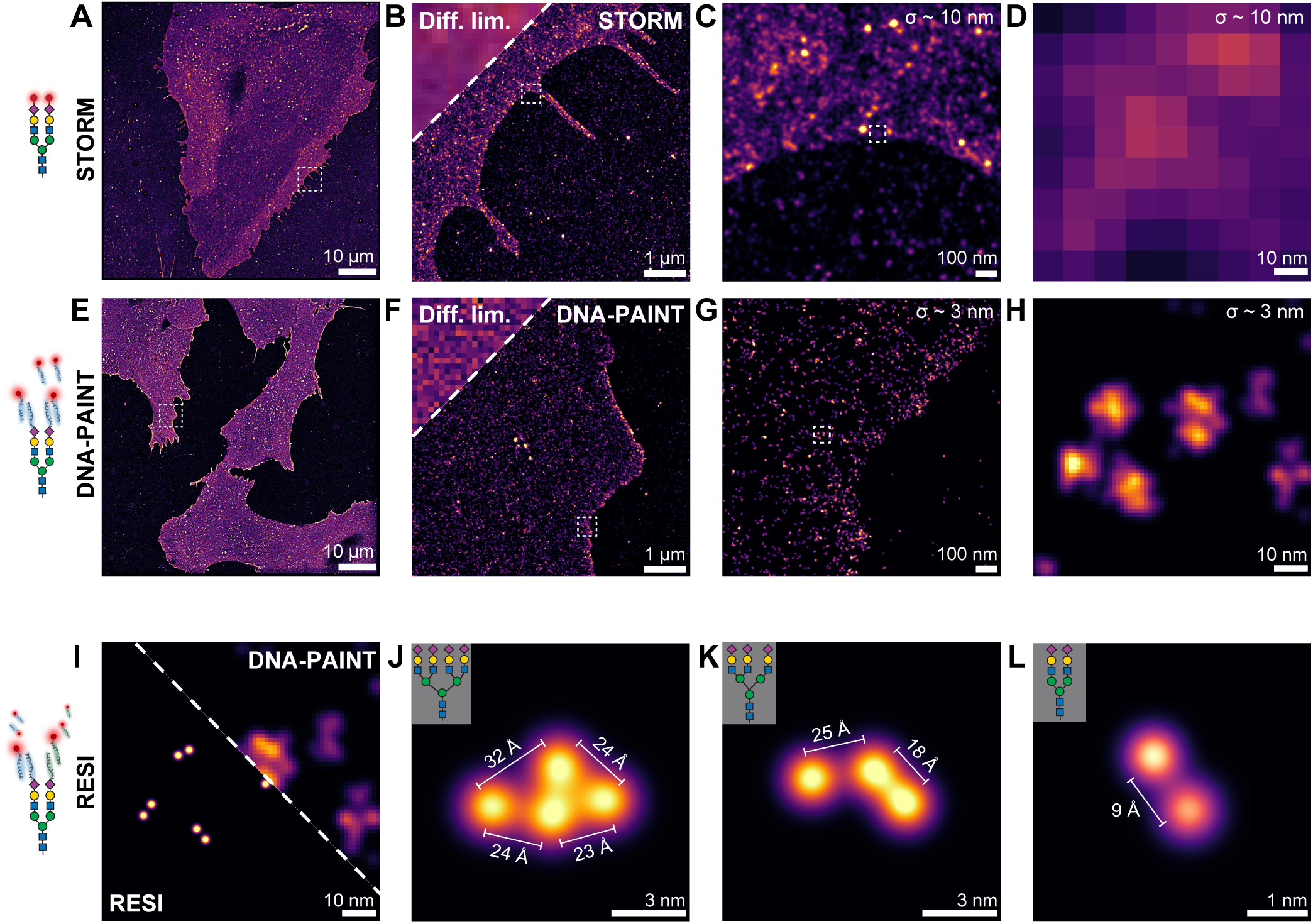
Visualization of cell surface sialic acids with Ångström-resolution. (**A**) Overview (d)STORM image of sialic acids on HMECs treated with ManNAz and labelled with A647-DBCO. (**B**) A side-by-side comparison of the diffraction-limited (upper left corner) and (d)STORM images shows improvement in resolution. (**C**) Zoomed-in (d)STORM image showing sialic acids on the cell surface. (**D**) Close-up view illustrating the limitations of (d)STORM in resolving individual sugar molecules. (**E**) Overview DNA-PAINT image of sialic acids on HMECs treated with ManNAz and labeled with a single DBCO-modified DNA sequence. (**F**) A side-by-side comparison of diffraction-limited (upper left corner) and DNA-PAINT images shows improved resolution. (**G**) Zoomed-in DNA-PAINT image showing sialic acids on the cell surface. (**H**) A close-up DNA-PAINT image showing approximately a 3-fold improvement in resolution compared to (d)STORM, though individual glycans remain unresolved. (**I**) Side-by-side comparison of RESI (left) and DNA-PAINT (right) capabilities in resolving sialic acids. Only RESI, but not DNA-PAINT, allows the detection of sub-10-nm sugar-sugar distances. (**J**) Tetra-antennary glycan on the cell surface. (**K**) Tri-antennary glycan on the cell surface. (**L**) Bi-antennary glycan capped with sialic acid residues that are 9 Å apart.

We then proceeded to HMECs labeled with DBCO-ssDNA using six different DNA sequences (see Methods). **Fig. 2E** shows a DNA-PAINT image encompassing several HMECs within a 100 x 100 μm^2^ field-of-view (FOV). **Fig. 2F-H** show successive zoom-ins. At ∼3 nm localization precision (∼7 nm resolution), DNA-PAINT offers approximately a 3-fold resolution increase compared to STORM (**Fig. 2F**) but still fails to resolve individual glycans (**Fig. 2H**), not to mention individual sugars within glycans. Only RESI (**Fig. 2 I-L**) at up to 3 Å localization precision allows us to resolve the molecular details of the glycocalyx, unveiling the spatial distribution and structure of single glycans and most excitingly their constituent sugars. For example, we resolve isolated clusters of single sugars that are compatible with bi-, tri-, and tetra-antennary glycans (**Fig. 2J-L**). Strikingly, as shown in **Fig. 2L**, we can resolve distances down to 9 Å between two single sugar residues in a glycan.

This demonstrates for the first time optical Ångström resolution in the native cellular context, extending fluorescence microscopy more than 250-fold over the diffraction limit of light while maintaining its main advantages in terms of low invasiveness and molecular specificity. Our approach combining RESI and metabolic labeling allows us to characterize the spatial arrangement of individual sugar residues within the native glycocalyx at previously unattainable molecular resolution. This achievement also represents the first demonstration of Ångström-resolution imaging of a complex, spatially extended structure in a whole cell, advancing beyond *in vitro* demonstrations (*31, 37*).

### The molecular arrangement of glycan-building units on the cell surface

Subsequently, we focused on extracting detailed quantitative information from the single-sugar-resolved datasets. We imaged HMECs labeled with Ac_4_ManNAz (**Fig. 3A-C and Fig. S3**) and Ac_4_GalNAz (**Fig. 3D-F and Fig. S3**), respectively. **Fig. 3A, D** show an overview of the cells, while **Fig. 3B, E**, and **Fig. 3C, F** show successive zoom-ins down to the scale of individual sugar molecules.

**Fig. 3.**
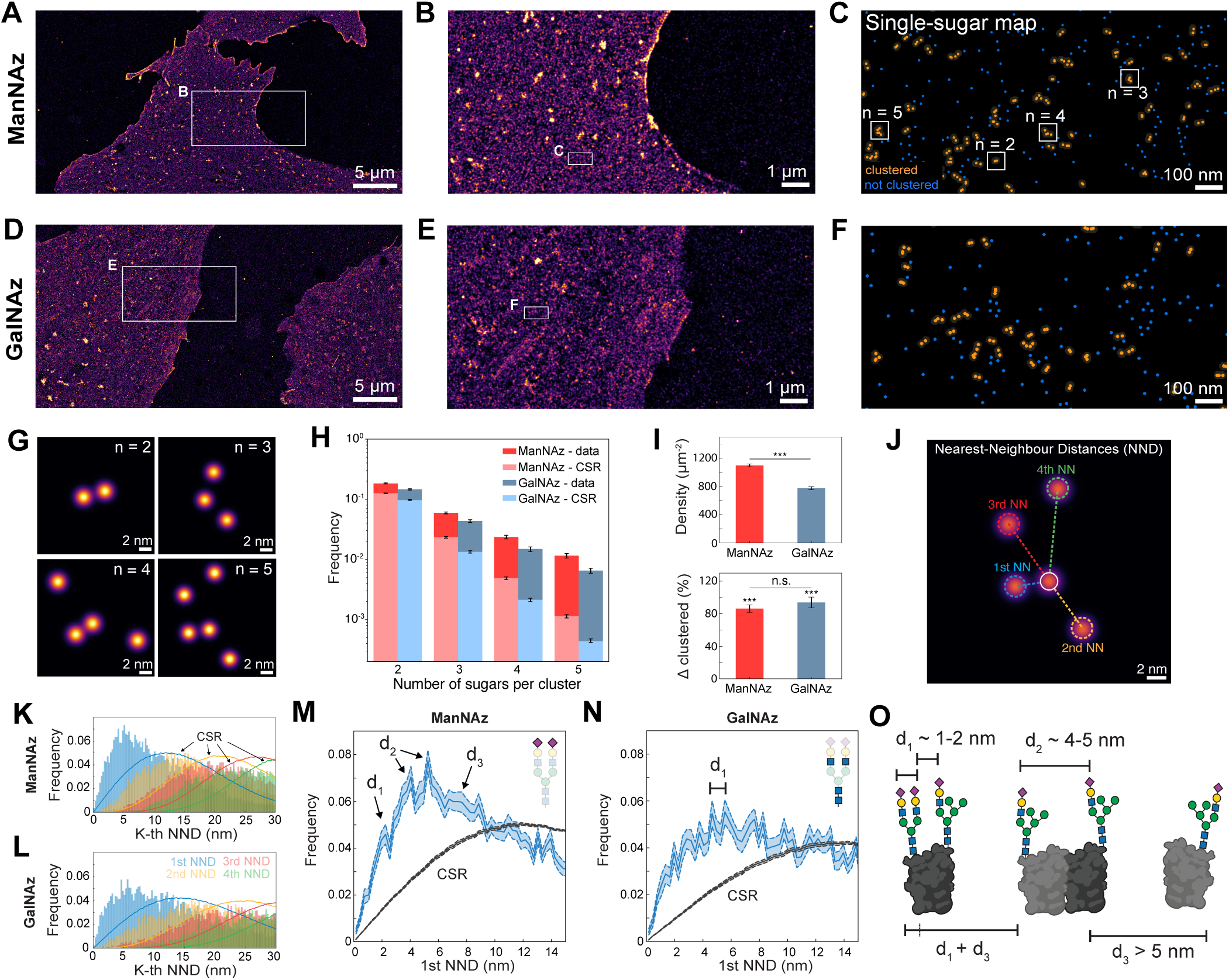
Identification of distinct nanoscale glycan fingerprints. (**A**) HMECs treated with ManNAz, targeting sialic acids, and labeled with six orthogonal DBCO-modified DNA sequences. (**B**) RESI zoom-in showing sialic acid residues on the cell surface. (**C**) Single sugars are clustered using DBSCAN. (**D**) HMECs treated with GalNAz, targeting LacNAc residues, and labeled with six orthogonal DBCO-modified DNA sequences. (**E**) RESI zoom-in showing LacNAc residues on the cell surface. (**F**) Representative clusters of LacNAc residues. (**G**) Single sugars visualized within individual clusters. (**H**) Frequency of clusters with five or fewer sugars compared to Complete Spatial Randomness (CSR). (**I**) Sugar density on the cell surface (top) and relative variation of the amount of clustered sugars (Δ clustered) compared to CSR (bottom). (**J**) Visual representation of first (blue), second (yellow), third (red), and fourth (green) nearest sugar distances for a given localization (white). (**K**) First to fourth nearest-neighbor sialic acid distances (histograms) compared to complete spatial randomness (dashed line). First, second, third, and fourth nearest neighbor distances are shown in blue, yellow, red, and green, respectively. (**L**) First to fourth nearest neighbor LacNAc distances (histograms) compared to complete spatial randomness (solid line). The color scheme is identical to subpanel K. (**M**) First-nearest neighbor distances for sialic acids (blue) and CSR (grey). The dotted lines and the shaded area represent the confidence interval of 67% (1 SD). (**N**) First-nearest neighbor distances for LacNAc residues (blue) and CSR (grey). The dotted lines and the shaded area represent the confidence interval of 67% (1 SD). (**O**) The quantitative analysis of the data is compatible with sugars located on different glycans of the same protein and sugars on neighboring glycoproteins.

To analyze the spatial distribution and molecular-scale organization of glycans on the cell surface, we applied the DBSCAN clustering algorithm (*38*) using a conservative distance threshold of d = 10 nm. This threshold was chosen based on an estimated typical glycan size of approximately 5-10 nm (*23*). By setting this value, we aimed to distinguish direct interactions (i.e., sugars within the same glycan, occurring at distances < 10 nm) from non-direct interactions (i.e., sugars located on different glycans, occurring at distances > 10 nm). **Fig. 3C, F** show clusters of sugars (orange) and single, non-clustered sugars (blue). Clusters of up to 5 sugars are detected on the cell surface (**Fig. 3G**). Canonical glycan branching stops at n = 4, however, a higher number of branches has been reported (*39, 40*). Thus, these clusters likely correspond to a single glycan with up to 5 branches or to more than one glycan bound to a protein.

In order to quantify the overall clusterization of sugar positions, we picked N = 20 different areas of 0.5 μm^2^ with apparently homogeneous density across different cells and different FOVs. We then histogrammed the number of sugars per cluster and compared the clusterization of the data with a Complete Spatial Randomness (CSR) simulation at the same densities (**Fig. 3H**). Both Ac_4_ManNAz (red) and Ac_4_GalNAz (blue) show a significantly higher clusterization than the one expected for CSR. Interestingly, despite observing differences in the overall sugar densities, the relative clusterization increase from CSR is the same for both Ac_4_ManNAz and Ac_4_GalNAz (**Fig. 3I**), indicating that the detected clusterization is not a density-dependent effect. For both Ac_4_ManNAz- and Ac_4_GalNAz-mediated labeling, we observe average densities of approximately 1100 and 750 sugars per µm², respectively, indicating high unnatural sugar incorporation and DBCO-click labeling efficiency (**Fig. 3I**). These high densities are consistent with glycosylation being the most abundant post-translational modification (*41*) and sialic acids serving as the most abundant terminal sugar on human cell-surface glycans (*42*). Ac_4_ManNAz is processed via the sialic acid biosynthetic pathway, where it is converted into CMP-Neu5Az and incorporated at glycan termini. Ac_4_GalNAz enters the hexosamine biosynthetic pathway, where it is metabolized into UDP-GalNAz and UDP-GlcNAz and incorporated into the core structures of glycans, likely causing broader metabolization and lower cell-surface density compared to Ac_4_ManNAz.

The cell surface is a densely packed environment, where proteins and sugar molecules are tightly arrayed (*43*). A single glycosylated protein, for example, can carry anywhere from a single glycan chain (e.g., a glycosylated membrane protein) to hundreds of glycan chains (in the case of mucins) (*44*). In order to further investigate the spatial arrangements within single-glycan length scales (up to 10 nm) we applied a Nearest-Neighbour Distance (NND) analysis (**Fig. 3J**). Briefly, the distances to the K^th^ NN were calculated for each sugar up to K = 4 and the distances were then histogrammed to obtain the NND distributions for both ManNAz (**Fig. 3K and Fig. S4**) and GalNAz (**Fig. 3L and Fig. S4**). All K^th^ NND distributions show a clear deviation from the CSR simulations (solid lines) at the same densities, indicating distinct molecular patterns at the sub-10 nm scale.

A closer inspection of the 1^st^ NND distributions shows distinct peaks at approximately 2, 4, and 6 nm for sialic acids (**Fig. 3M**) and sub-6 nm peaks spaced by approx. 1 nm for Ac_4_GalNAz-labeled cells (**Fig. 3N**). Given that the average diameter of an integral membrane protein is approximately 5 nm (*45*), the distances of ManNAz are fully consistent with i) sialic acids capping the same glycan (**Fig. S5**), ii) sialic acids located on different glycans of the same protein, and iii) sialic acids on neighboring glycoproteins, respectively (**Fig. 3O**). On the other hand, while sialic acids are located at the terminal positions of glycan chains (**Fig. 3M**), LacNAc residues are more densely packed and positioned closer together due to their presence within the glycan core (**Fig. 3N**). Here, distinct peaks at distances of approximately 1, 2, and 3 nm are visible. These are consistent with *in silico* distance measurements, where a set of biologically relevant glycan structures was surveyed (**Fig. S5**).

## Conclusions

Our work provides the first visualization of the molecular structure of the glycocalyx in its native cellular context. This novel capability is poised to have far-reaching implications for functional glycobiology and cell biology in general. For the first time, glycocalyx components can now be studied in their native environment with spatial resolution down to a single sugar unit, which has been considered one of the central desiderata of glycobiology (*46*).

A persistent challenge in glycobiology has been the lack of a defined composition for a healthy glycocalyx: its aberrant states still rely on the same fundamental building blocks—there are no inherently “diseased” glycocalyx components per se. This complicates drug discovery in this field, as both excessive and insufficient levels of certain glycocalyx constituents can lead to severe side effects (*47*). For example: hypersialylation can fuel tumour growth and enable immune evasion (*48*), while hyposialylation may trigger autoimmune diseases (*49*). By enabling Ångström-resolution mapping of glycans, RESI paves the way to unveil what a healthy glycocalyx looks like at the molecular level and observe how drugs alter its organization. This opens the door to the rational development of highly specific glycocalyx-targeting therapies, transforming how we approach drug discovery in this field.

Understanding glycan-dependent processes such as glycan-receptor interactions requires imaging methods that can reveal glycans and interacting proteins with molecular resolution in a multiplexed fashion. By combining Ångström-resolution imaging of glycans with expanded DNA barcodes (*50*), one could simultaneously visualize membrane receptor proteins, providing new insights into their spatial organization and interactions. RESI is particularly well suited for multiplexing as it decouples species identification from fluorophore wavelength, instead encoding molecular identity in DNA sequences used for imaging (*51*).

From an imaging technology point of view, we have shown that combining RESI with metabolic labeling unlocks the full potential of Ångström-resolution fluorescence microscopy, demonstrating – for the first time – a resolution below 1 nm in a native cellular environment while maintaining fields-of-view of 100 x 100 μm^2^ comprising several cells per image acquisition. We envision that click-chemistry approaches using unnatural amino acids (*52*) could extend the Ångström-resolution achieved in glycans to proteins and other types of biomolecules.

From a methodological perspective, our work underscores the urgent need for rapid and extensive advancements in labeling glycocalyx constituents. While the frontier in glycocalyx research was once defined by the limitations of optical resolution, we have now surpassed those constraints, achieving single-sugar resolutions. However, the field continues to face a significant gap in tools capable of effectively targeting and labeling relevant glycocalyx components.

Looking ahead, expanding the current repertoire of metabolic labeling tools beyond those targeting LacNAc and sialic acids will be crucial, alongside driving the development of novel strategies for glycan labeling. RESI could significantly accelerate these efforts by enabling the precise analysis of labeling performance and providing insights into how incorporation efficiency is influenced by changes in the biosynthetic machinery of cells. With advancements in glycan labeling techniques, RESI is poised to resolve complete glycan structures, paving the way for glycoproteomics using light microscopy.

Finally, given the growing importance of cancer glycoimmunology and clinical glycobiology, our approach could significantly deepen our understanding of the functional role of glycosylation in cancer biology. Precise mapping of changes in glycan organization, branching, and density during cancer progression could add a new axis to the clinical analysis of glycosylation. Accordingly, our method could enable a completely new understanding of the role of cancer glycosylation in immune system regulation and immune evasion (*53, 54*). This will be of paramount importance for the identification of new diagnostic markers and targets in cancer immunotherapy, where glycosylation is already recognized as a central regulator. In conclusion, our findings will not only elucidate unknown areas of fundamental glycoscience but could also directly improve diagnosis and therapy.

## Materials and Methods

### Cell culture

All cells were cultured in T75 flasks (Corning BV) in a humidified atmosphere at 37°C with 5% CO_2_. HMECs were cultured in MDCB-131 medium with 1 % Glutamax, 10 % fetal bovine serum (FBS), 10 ng/mL hEGF (all Thermo Fisher Scientific), 1 μg/mL hydrocortisone, and 1 % of a penicillin-streptomycin solution containing 10,000 U/mL penicillin and 10 mg/L streptomycin (both Sigma-Aldrich). For imaging, cells were seeded on Lab-Tek II Chambered Coverglass (Thermo Fisher Scientific).

### DNA-PAINT sequences

Six orthogonal DNA sequences modified with aza-dibenzocyclooctyne (DBCO) at their 5’ ends were used to label the azido-sugars. The docking strand sequences used were 5xR1 (TCCTCCTCCTCCTCCTCCT), 5xR2 (ACCACCACCACCACCACCA), 7xR3 (CTCTCTCTCTCTCTCTCTC), and 7xR4 (ACACACACACACACACACA), 5xR5 (CTTCTTCTTCTTCTTCTTC), and 5x R6 (AACAACAACAACAACAACAA). Their respective imagers were R1 (AGGAGGA-Cy3B), R2 (GGTGGT-Cy3B), R3 (GAGAGAG-Cy3B), R4 (TGTGTGT-Cy3B), R5 (GAAGAAG-Cy3B), and R6 (TG TTG TT-Cy3B). All sequences were speed-optimized (*56*). The docking strand and imager sequences were purchased from Metabion.

### DNA-PAINT imaging buffer

1× PBS, 1 mM EDTA, 500 mM NaCl pH 7.4, 0.02% Tween, optionally supplemented with 1× Trolox, 1× PCA and 1× PCD. Tween-20 (no. P9416-50ML), protocatechuate 3,4-dioxygenase pseudomonas (PCD, no. P8279), 3,4-dihydroxybenzoic acid (PCA, no. 37580-25G-F), 1× PBS (pH 7.2, no. 20012-019) and (±)-6-hydroxy-2,5,7,8-tetra-methylchromane-2-carboxylic acid (Trolox, no. 238813-5G) were ordered from Sigma-Aldrich. EDTA (no. AM9260G), and 1× PBS (pH 7.2, no. 20012-019) were purchased from Thermo Fisher Scientific.

### Metabolic incorporation of Ac4GalNAz/Ac4ManNAz, click of docking strands, and fixation

A few hours after seeding, the cells were supplemented with 50 μM of Ac4ManNAz (Thermo Fisher Scientific) or Ac4GalNAz (Thermo Fisher Scientific) in cell culture media. After 72 hours of incubation, when cells had reached approximately 60% confluency, they were treated with 50 μM of each of the six docking DNA strands in culture media for 2 hours at 37°C to allow for the click reaction between the docking DNA sequences and the azido sugars. Following this incubation, the cells were fixed with 4% paraformaldehyde (Sigma-Aldrich) in DPBS (Life Technologies) for 20 minutes at room temperature. The cells were then permeabilized using 0.1% Triton X-100 (Thermo Fisher Scientific) in DPBS for 10 minutes at room temperature. To ensure thorough removal of unreacted reagents and minimize background, cells were washed three times with DPBS at each step, including before and after the click reaction, fixation, and permeabilization procedures.

### Preparation of STORM samples

For STORM imaging, samples were prepared using the same protocol as described above, with one modification: instead of docking DNA strands, 50 μM of DBCO-647 (Jena Bioscience) was used for the click reaction.

### STORM imaging

A reducing oxygen scavenging buffer that induces blinking of single fluorophores was employed according to the literature (*57*). The STORM buffer consisted of 2 μL/mL catalase (Sigma-Aldrich), 10% (w/v) glucose (BD Biosciences), 100 mM Tris-HCl (Thermo Fisher Scientific), 560 μg/mL glucose oxidase (Sigma-Aldrich), and 20 mM cysteamine (Sigma-Aldrich). The PBS in which the fixed cells were stored was replaced by the blinking buffer. First, diffraction-limited (DL) imaging was performed with low-intensity illumination of 1 W/cm^2^. Then, the laser power was increased to ∼1.2 kW/cm^2^. Image acquisition was started after a short delay to ensure that most fluorophores were shelved into a dark state. The exposure time was 50 ms, and 40,000 frames were obtained for sialic acid imaging.

### RESI imaging

Gold nanoparticles (Cytodiagnostics, no. G-90-100) were diluted 1:2 in PBS and incubated for 10 min at RT and the sample was washed two times with PBS to remove unbound gold. First, the imager solution (see Supplementary Table 1) in DNA-PAINT imaging buffer for the first round was incubated for 2 min and then replaced with fresh imager, after which the first acquisition round was started. The sample was washed with at least 2 mL of PBS between imaging rounds until no residual signal from the previous imager solution was detected. Then, the next imager solution was introduced. RESI imaging was conducted via six subsequent DNA-PAINT imaging rounds with only one of the imagers in each round. The optimal imager concentration required to achieve sparse blinking can vary. In this study, we utilized concentrations up to 125 pM. These concentrations were adjusted to ensure that blinking events were frequent enough yet sparse enough to maintain high DNA-PAINT resolution. The optimal concentration for each dataset was determined through visual assessment of the blinking and remained consistent across the corresponding set to allow for meaningful comparisons. In each field of view, 40,000 frames with 100 ms exposure time per frame were acquired. A laser power of 35 mW (560 nm laser, measured after the objective) was used, corresponding to a power density of ∼175 W/cm^2^. Further details on imaging parameters are provided in Supplementary Table 1.

### Microscopy setup

Fluorescence imaging was carried out on an inverted microscope (Nikon Instruments, Eclipse Ti2) with the Perfect Focus System, applying an objective-type TIRF configuration equipped with an oil-immersion objective (Nikon Instruments, Apo SR TIRF×100, NA 1.49, Oil). A 560-nm laser (MPB Communications, 1 W) was used for excitation and coupled into the microscope via a Nikon manual TIRF module. The laser beam was passed through a cleanup filter (Chroma Technology, ZET561/10) and coupled into the microscope objective using a beam splitter (Chroma Technology, ZT561rdc). Fluorescence was spectrally filtered with an emission filter (Chroma Technology, ET600/50m, and ET575lp) and imaged on an sCMOS camera (Hamamatsu Fusion BT) without further magnification, resulting in an effective pixel size of 130 nm after 2×2 binning. TIR illumination was used for all measurements. The central 1152×1152 pixels (576×576 after binning) of the camera were used as the region of interest. The scan mode of the camera was set to “ultra quiet scan” (readout noise = 0.7 e-r.m.s., 80 μs readout time per line). Raw microscopy data was acquired using μManager (Version 2.0.1) (*58*).

### Single-molecule localization analysis

Raw fluorescence data were reconstructed using the Picasso software package (*35*) (the latest version is available at https://github.com/jungmannlab/picasso). Drift correction was performed using the AIM algorithm (*59*) with gold nanoparticles as fiducials for all experiments. The six channels were aligned through cross-correlation of the fiducial gold nanoparticles using Picasso. DNA-PAINT images displayed in Fig. 2 are obtained by merging all six RESI acquisition channels after alignment.

### RESI analysis

RESI analysis was carried out as described previously (*31*). Briefly, the localizations in each of the six channels were clustered using the custom clustering algorithm described previously. The clustering algorithm uses two input parameters: radius *r*, which sets the final size of the clusters and defines a circular environment around each localization, and the minimal number of localizations, *nmin*, representing a lower threshold for the number of DNA-PAINT localizations in any cluster. We used a radius *r* = 7.15 nm and *nmin* = 10 localizations. The radius value *r* is chosen to be approx. 2.35 times our localization precision which yields optimal results in terms of false detections and the minimum number of *nmin* = 10 localizations ensures the distinction of repetitively sampled binding sites (single sugars) from unspecific binding events of the imagers. Clusters of localizations are further analyzed in the time domain (time traces) and unspecific binding events are filtered out. Finally, the center of each cluster is calculated and the cluster centers of all six channels are merged to produce the RESI image.

### Quantitative analysis of RESI data

Downstream analysis (DBSCAN and Nearest Neighbour Distances) of the RESI data was performed using custom-written Python scripts based on Numpy functions (*60*) and Scipy (*61*) implementations of DBSCAN (*38*) and KD-tree Nearest Neighbour search (*62*).

### Computational sugar-sugar distance analysis

To measure sugar-sugar distances within glycans, we performed structural modeling and analysis using Chimera (*63*) version 1.18. Glycan conformations were optimized via Gasteiger energy minimization to ensure accurate modeling. Distances were then computationally determined by measuring the spatial separation between the carbon atoms nearest to the anchor points of the experimental labels. This approach provided theoretical distance estimations, aligning the computational model with the expected experimental labeling positions.

## Data availability

All data and analysis codes are available upon request.

## Author contributions

L.A.M. and K.A. built the optical systems, designed and performed all the experiments, and analyzed the data. I.P., M.H., and L.H. provided support with experiments. S.F. and H.G. provided support with data analysis. L.A.M., K.A., L.M., and R.J. interpreted data and wrote the manuscript with input from all authors. L.M. and R.J. conceived and supervised the project. L.A.M and K.A. contributed equally. Co-first authors L.A.M. and K.A. and co-corresponding authors R.J. and L.M. equally share the respective contribution, and each may list themselves first/last in order on their CVs.

## Funding

K.A., S.F., and L.M. gratefully acknowledge financial support from the Else-Kröner-Fresenius-Stiftung (grant ID 2020_EKEA.91), the German Research Foundation (DFG, grant ID 529257351), and the Wilhelm-Sander-Stiftung (grant ID 2023.025.1), as well as by the Max Planck Society. L.A.M. acknowledges a postdoctoral fellowship from the European Union’s Horizon 20212022 research and innovation program under Marie Skłodowska-Curie grant agreement no. 101065980. I.P. and M.H. acknowledge support from the International Max Planck Research School for Molecules of Life (IMPRS-ML). This research was funded in part by the European Research Council through an ERC Consolidator Grant (ReceptorPAINT, grant agreement no. 101003275), the Danish National Research Foundation (Centre for Cellular Signal Patterns, DNRF135), the Volkswagen Foundation through the initiative ‘Life?—A Fresh Scientific Approach to the Basic Principles of Life’ (grant no. 98198), and the Max Planck Foundation.

## Competing interests

The authors declare no competing financial interest.

## Supplementary Information

**Fig. S1.**
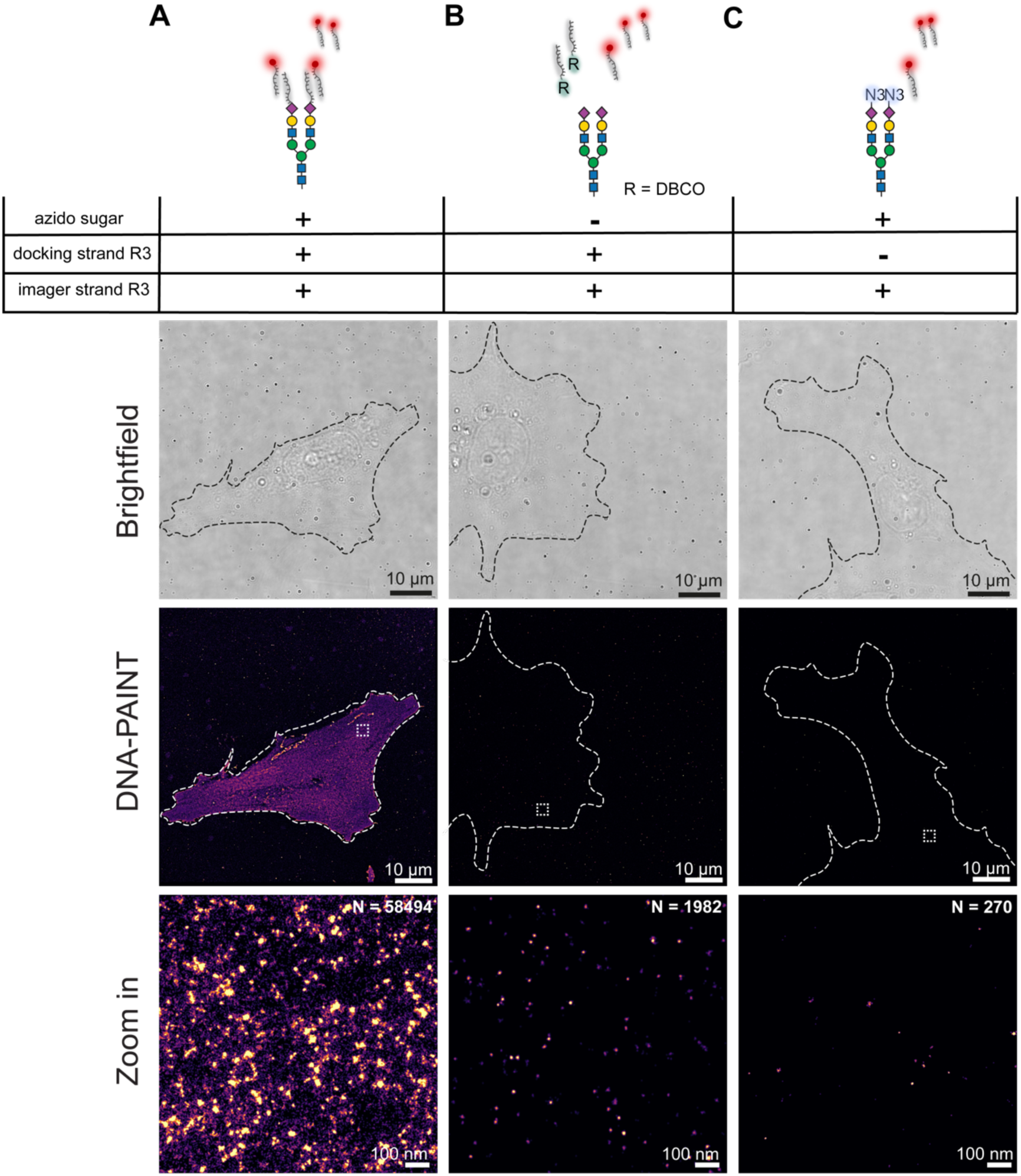
Validation of the labeling workflow. (**A**) Specificity of sialic acid labeling is demonstrated in comparison to controls: (**B**) samples prepared without azido sugars and (**C**) samples prepared without DNA docking strands. The number of localizations (N) is indicated in the upper-right corner of the zoomed-in panels, highlighting the labeling specificity.

**Fig. S2.**
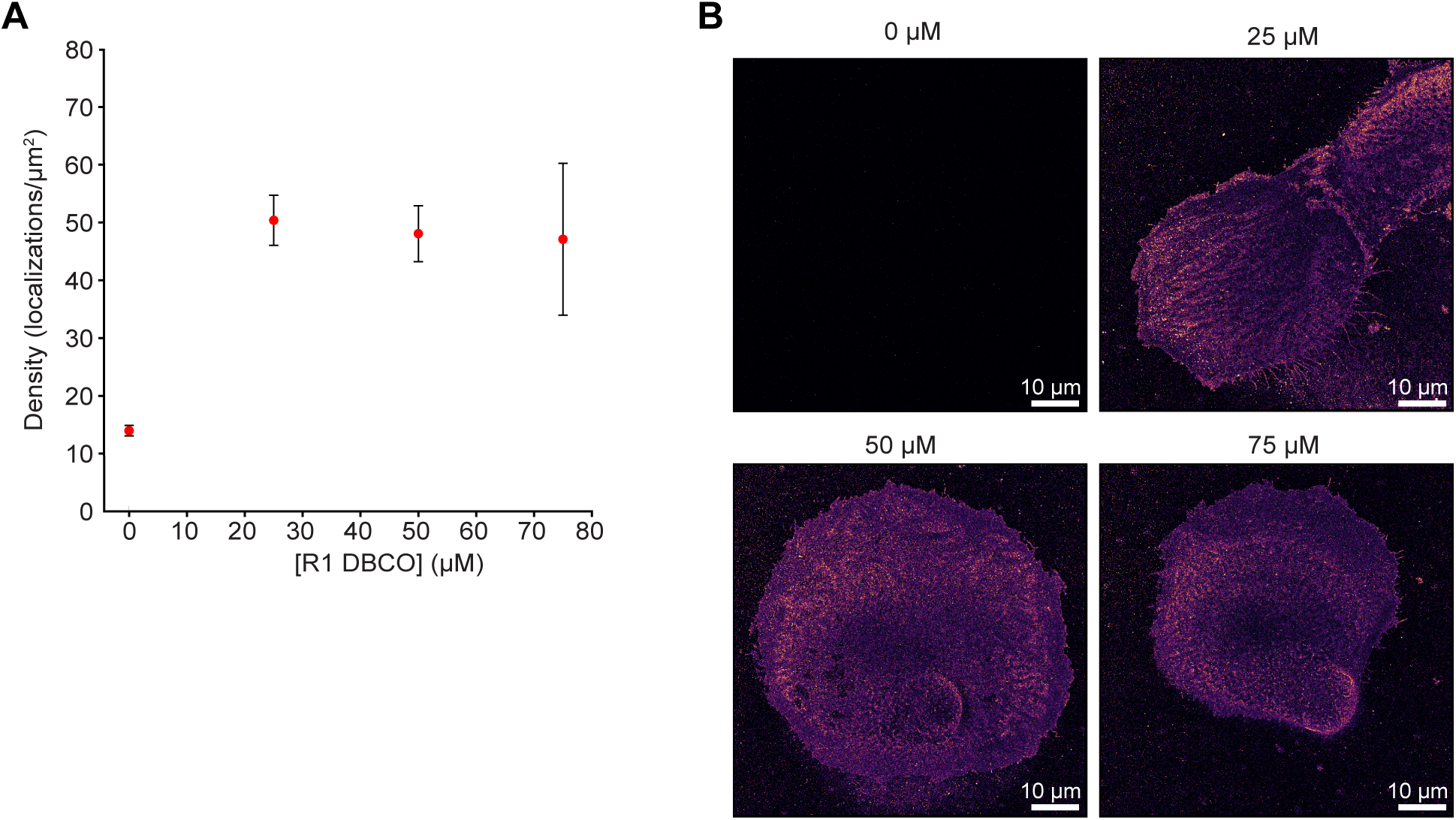
Saturation of azido binding sites. (**A**) Graphical representation of binding site saturation as a function of DBCO-R1 concentration. (**B**) Representative images demonstrating binding site saturation. Error bars indicate the standard error of the mean (SEM). N=6.

**Fig. S3.**
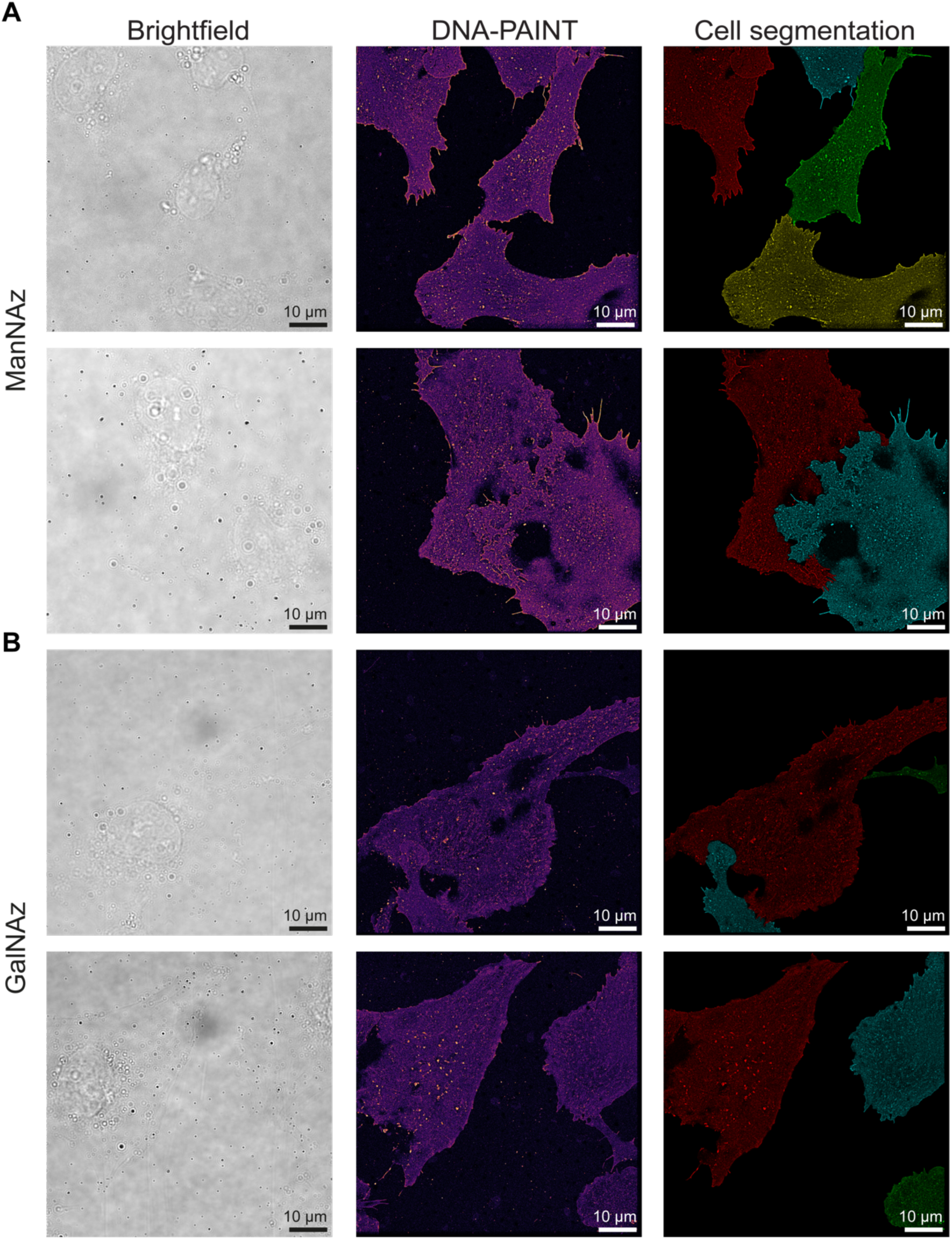
Extended data. (**A**) HMECs labeled with ManNAz. (**B**) HMECs labeled with GalNAz. The left panels show brightfield images, the middle panels display DNA-PAINT overviews, and the right panels show segmented cells, with each cell depicted in a different color.

**Fig. S4.**
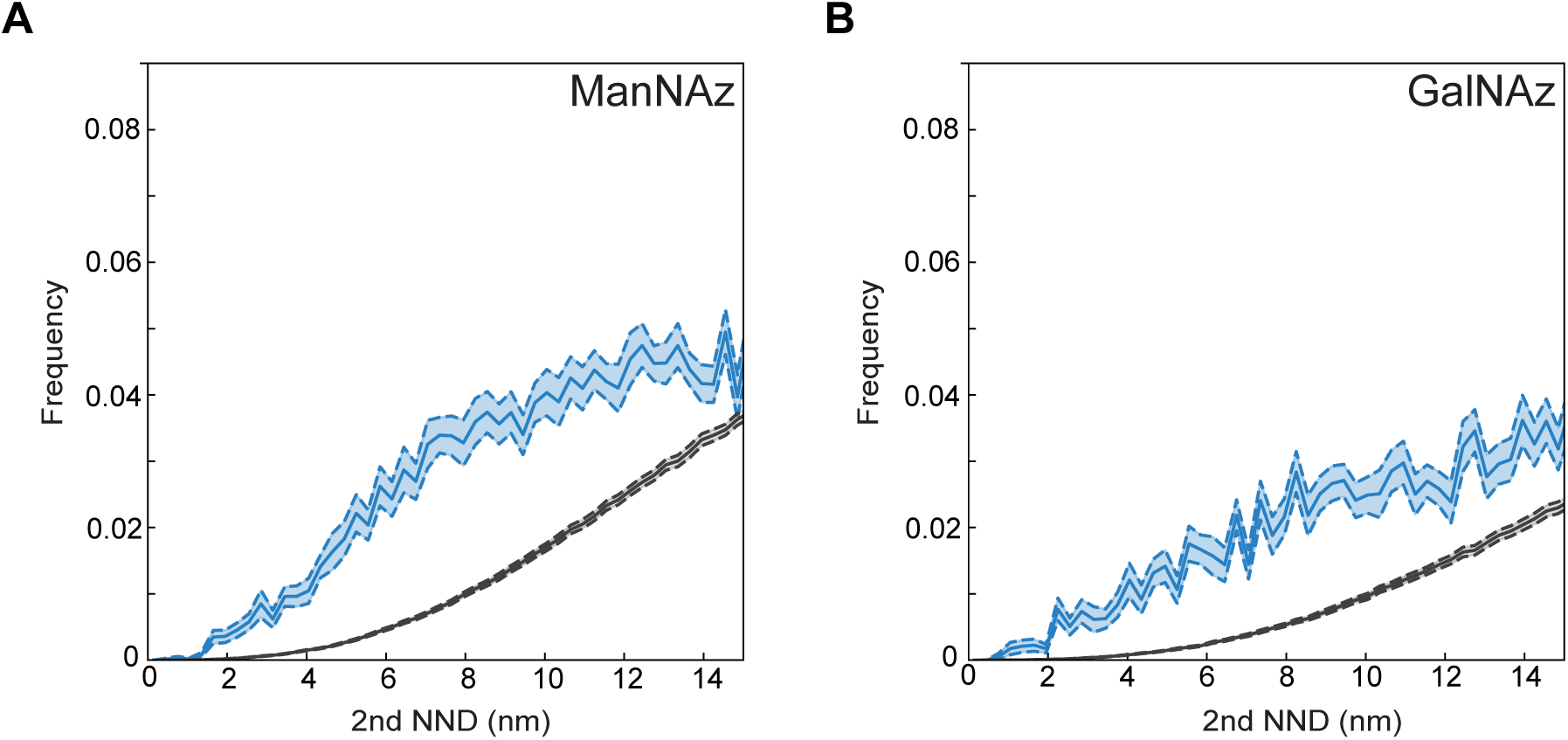
Second-nearest neighbor distances. (**A**) HMECs labeled with ManNAz. (**B**) HMECs labeled with GalNAz.

**Fig. S5.**
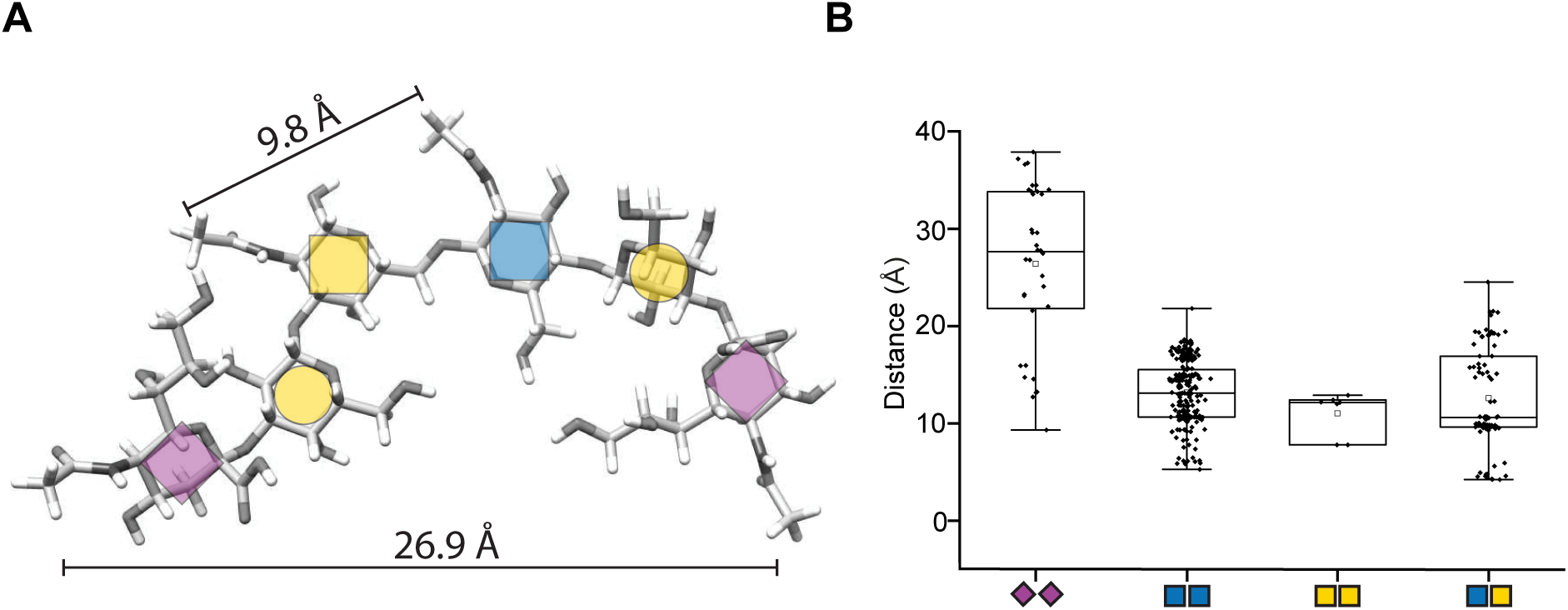
Sugar-sugar distances within a glycan. (**A**) Theoretical sugar-sugar distances measured within a glycan. (**B**) Sugar-sugar distances between sialic acids (labeled with ManNAz) and GlcNAc/GalNAc (labeled with GalNAz) measured across glycan structures listed on GlycoShape (*64*).

### Supplementary Tables

**Supplementary Table 1.** Acquisition parameters.

| Data | Round | Exposure (ms) | Frames | Power (at the objective, mW) | Imagers | Buffer | Cluster Radius xy (nm) | Cluster N<sub>min</sub> |
| --- | --- | --- | --- | --- | --- | --- | --- | --- |
| (d)STORM<br>Fig. 2 (A-D) | Round 1 | 50 | 40 000 | 200 | Alexa 647 | STORM buffer | - | - |
| RESI<br>Figure 2 (I-L),<br>Figure 3 | Round 1 | 100 | 40 000 | 35 | 125 pM R1 | DNA-PAINT buffer | 7.15 | 10 |
|  | Round 2 |  | 40 000 |  | 50 pM R2 |  |  |  |
|  | Round 3 |  | 40 000 |  | 50 pM R3 |  |  |  |
|  | Round 4 |  | 40 000 |  | 5 pM R4 |  |  |  |
|  | Round 5 |  | 40 000 |  | 125 pM R5 |  |  |  |
|  | Round 6 |  | 40 000 |  | 125 pM R6 |  |  |  |
| Fig S1 | Round 1 | 100 | 40 000 | 35 | 125 pM R3 | DNA-PAINT buffer | - | - |
| Fig S2 | Round 1 | 100 | 20 000 | 35 | 250 pM R1 | DNA-PAINT buffer | - | - |

**Supplementary Table 2.** Docking sequences.

| Docking strand name | Sequence |
| --- | --- |
| 5xR1 | TCCTCCTCCTCCTCCT |
| 5xR2 | ACCACCACCACCACCA |
| 7xR3 | CTCTCTCTCTCTCTC |
| 7xR4 | ACACACACACACACACA |
| 5xR5 | CTTCTTCTTCTTCTTC |
| 5xR6 | AACAACAACAACAACAA |

**Supplementary Table 3.** Imager sequences.

| Imager name | Sequence | 3'-mod |
| --- | --- | --- |
| R1 | AGGAGGA | Cy3b |
| R2 | GGTGGT | Cy3b |
| R3 | GAGAGAG | Cy3b |
| R4 | TGTGTGT | Cy3b |
| R5 | GAAGAAG | Cy3b |
| R6 | TGTTGTT | Cy3b |

## References

1. R. A. Flynn et al., Small RNAs are modified with N-glycans and displayed on the surface of living cells. Cell 184, 3109–3124 e3122 (2021).

2. J. M. Tarbell, L. M. Cancel, The glycocalyx and its significance in human medicine. J Intern Med 280, 97–113 (2016).

3. L. Möckl, The Emerging Role of the Mammalian Glycocalyx in Functional Membrane Organization and Immune System Regulation. Front Cell Dev Biol 8, 253 (2020).

4. P. R. Crocker, J. C. Paulson, A. Varki, Siglecs and their roles in the immune system. Nat Rev Immunol 7, 255–266 (2007).

5. B. A. H. Smith, C. R. Bertozzi, The clinical impact of glycobiology: targeting selectins, Siglecs and mammalian glycans. Nat Rev Drug Discov 20, 217–243 (2021).

6. H. H. Lipowsky, The endothelial glycocalyx as a barrier to leukocyte adhesion and its mediation by extracellular proteases. Ann Biomed Eng 40, 840–848 (2012).

7. M. J. Paszek et al., The cancer glycocalyx mechanically primes integrin-mediated growth and survival. Nature 511, 319–325 (2014).

8. I. Bagdonaite et al., Glycoproteomics. Nature Reviews Methods Primers 2, 48 (2022).

9. X. Wu et al., Imaging single glycans. Nature 582, 375–378 (2020).

10. K. P. Arkill et al., 3D reconstruction of the glycocalyx structure in mammalian capillaries using electron tomography. Microcirculation 19, 343–351 (2012).

11. L. Möckl et al., Quantitative Super-Resolution Microscopy of the Mammalian Glycocalyx. Dev Cell 50, 57–72 e56 (2019).

12. C. T. McDowell, X. Lu, A. S. Mehta, P. M. Angel, R. R. Drake, Applications and continued evolution of glycan imaging mass spectrometry. Mass Spectrom Rev 42, 674–705 (2023).

13. K. Anggara et al., Direct observation of glycans bonded to proteins and lipids at the single-molecule level. Science 382, 219–223 (2023).

14. E. E. Ebong, F. P. Macaluso, D. C. Spray, J. M. Tarbell, Imaging the endothelial glycocalyx in vitro by rapid freezing/freeze substitution transmission electron microscopy. Arterioscler Thromb Vasc Biol 31, 1908–1915 (2011).

15. J. Hegermann, H. Lunsdorf, M. Ochs, H. Haller, Visualization of the glomerular endothelial glycocalyx by electron microscopy using cationic colloidal thorium dioxide. Histochem Cell Biol 145, 41–51 (2016).

16. L. Chevalier et al., Electron microscopy approach for the visualization of the epithelial and endothelial glycocalyx. Morphologie 101, 55–63 (2017).

17. K. Kappler, T. Hennet, Emergence and significance of carbohydrate-specific antibodies. Genes Immun 21, 224–239 (2020).

18. P. Stanley, Genetics of glycosylation in mammalian development and disease. Nat Rev Genet 25, 715–729 (2024).

19. R. D. Cummings et al., in Essentials of Glycobiology, A. Varki et al., Eds. (Cold Spring Harbor (NY), 2022), pp. 645–662.

20. J. M. Baskin et al., Copper-free click chemistry for dynamic in vivo imaging. Proc Natl Acad Sci U S A 104, 16793–16797 (2007).

21. S. T. Laughlin, J. M. Baskin, S. L. Amacher, C. R. Bertozzi, In vivo imaging of membrane-associated glycans in developing zebrafish. Science 320, 664–667 (2008).

22. S. Letschert et al., Super-resolution imaging of plasma membrane glycans. Angew Chem Int Ed Engl 53, 10921–10924 (2014).

23. C. Reily, T. J. Stewart, M. B. Renfrow, J. Novak, Glycosylation in health and disease. Nat Rev Nephrol 15, 346–366 (2019).

24. M. Aebi, R. Bernasconi, S. Clerc, M. Molinari, N-glycan structures: recognition and processing in the ER. Trends Biochem Sci 35, 74–82 (2010).

25. M. J. Rust, M. Bates, X. Zhuang, Sub-diffraction-limit imaging by stochastic optical reconstruction microscopy (STORM). Nat Methods 3, 793–795 (2006).

26. M. Heilemann et al., Subdiffraction-resolution fluorescence imaging with conventional fluorescent probes. Angew Chem Int Ed Engl 47, 6172–6176 (2008).

27. P. Mateos-Gil, S. Letschert, S. Doose, M. Sauer, Super-Resolution Imaging of Plasma Membrane Proteins with Click Chemistry. Front Cell Dev Biol 4, 98 (2016).

28. R. Riera et al., Single-molecule imaging of glycan-lectin interactions on cells with Glyco-PAINT. Nat Chem Biol 17, 1281–1288 (2021).

29. D. A. Helmerich et al., Photoswitching fingerprint analysis bypasses the 10-nm resolution barrier. Nat Methods 19, 986–994 (2022).

30. K. Almahayni, M. Spiekermann, L. Mockl, Fluorophores’ talk turns them dark. Nat Methods 19, 932–933 (2022).

31. S. C. M. Reinhardt et al., Angstrom-resolution fluorescence microscopy. Nature 617, 711–716 (2023).

32. S. T. Laughlin, C. R. Bertozzi, Metabolic labeling of glycans with azido sugars and subsequent glycan-profiling and visualization via Staudinger ligation. Nat Protoc 2, 2930–2944 (2007).

33. 33. S. L. Scinto, et al., Bioorthogonal chemistry. *Nat Rev Methods Primers* 1, (2021).

34. R. Jungmann et al., Single-molecule kinetics and super-resolution microscopy by fluorescence imaging of transient binding on DNA origami. Nano Lett 10, 4756–4761 (2010).

35. J. Schnitzbauer, M. T. Strauss, T. Schlichthaerle, F. Schueder, R. Jungmann, Super-resolution microscopy with DNA-PAINT. Nat Protoc 12, 1198–1228 (2017).

36. D. Axelrod, N. L. Thompson, T. P. Burghardt, Total internal reflection fluorescent microscopy. J Microsc 129, 19–28 (1983).

37. S. J. Sahl et al., Direct optical measurement of intramolecular distances with angstrom precision. Science 386, 180–187 (2024).

38. M. Ester, H.-P. Kriegel, J. Sander, X. Xu, paper presented at the Proceedings of the Second International Conference on Knowledge Discovery and Data Mining, Portland, Oregon, 1996.

39. N. Suzuki, T. Abe, K. Hanzawa, S. Natsuka, Toward robust N-glycomics of various tissue samples that may contain glycans with unknown or unexpected structures. Sci Rep 11, 6334 (2021).

40. Z. Sumer-Bayraktar et al., N-glycans modulate the function of human corticosteroid-binding globulin. Mol Cell Proteomics 10, M111 009100 (2011).

41. K. T. Schjoldager, Y. Narimatsu, H. J. Joshi, H. Clausen, Global view of human protein glycosylation pathways and functions. Nat Rev Mol Cell Biol 21, 729–749 (2020).

42. R. L. Schnaar, Glycobiology simplified: diverse roles of glycan recognition in inflammation. J Leukoc Biol 99, 825–838 (2016).

43. C. R. Shurer et al., Physical Principles of Membrane Shape Regulation by the Glycocalyx. Cell 177, 1757–1770 e1721 (2019).

44. G. Raba, A. S. Luis, Mucin utilization by gut microbiota: recent advances on characterization of key enzymes. Essays Biochem 67, 345–353 (2023).

45. J. Chin-Hun Kuo, J. G. Gandhi, R. N. Zia, M. J. Paszek, Physical biology of the cancer cell glycocalyx. Nat Phys 14, 658–669 (2018).

46. A. Lakshminarayanan, M. Richard, B. G. Davis, Studying glycobiology at the single-molecule level. Nature Reviews Chemistry 2, 148–159 (2018).

47. K. Almahayni, L. Mockl, Setting the stage for universal pharmacological targeting of the glycocalyx. Curr Top Membr 91, 61–88 (2023).

48. C. Dobie, D. Skropeta, Insights into the role of sialylation in cancer progression and metastasis. Br J Cancer 124, 76–90 (2021).

49. A. Bordron et al., Hyposialylation Must Be Considered to Develop Future Therapies in Autoimmune Diseases. Int J Mol Sci 22, (2021).

50. E. M. Unterauer et al., Spatial proteomics in neurons at single-protein resolution. Cell 187, 1785–1800 e1716 (2024).

51. R. Jungmann et al., Multiplexed 3D cellular super-resolution imaging with DNA-PAINT and Exchange-PAINT. Nat Methods 11, 313–318 (2014).

52. M. Budiarta, M. Streit, G. Beliu, Site-specific protein labeling strategies for super-resolution microscopy. Curr Opin Chem Biol 80, 102445 (2024).

53. J. E. Hudak, S. M. Canham, C. R. Bertozzi, Glycocalyx engineering reveals a Siglec-based mechanism for NK cell immunoevasion. Nat Chem Biol 10, 69–75 (2014).

54. B. S. Blaum et al., Structural basis for sialic acid-mediated self-recognition by complement factor H. Nat Chem Biol 11, 77–82 (2015).

55. S. Neelamegham et al., Updates to the Symbol Nomenclature for Glycans guidelines. Glycobiology 29, 620–624 (2019).

56. S. Strauss, R. Jungmann, Up to 100-fold speed-up and multiplexing in optimized DNA-PAINT. Nat Methods 17, 789–791 (2020).

57. A. R. Halpern, M. D. Howard, J. C. Vaughan, Point by Point: An Introductory Guide to Sample Preparation for Single-Molecule, Super-Resolution Fluorescence Microscopy. Curr Protoc Chem Biol 7, 103–120 (2015).

58. A. D. Edelstein et al., Advanced methods of microscope control using muManager software. J Biol Methods 1, (2014).

59. H. Ma, M. Chen, P. Nguyen, Y. Liu, Toward drift-free high-throughput nanoscopy through adaptive intersection maximization. Sci Adv 10, eadm7765 (2024).

60. C. R. Harris et al., Array programming with NumPy. Nature 585, 357–362 (2020).

61. P. Virtanen et al., SciPy 1.0: fundamental algorithms for scientific computing in Python. Nat Methods 17, 261–272 (2020).

62. J. L. Bentley, Multidimensional binary search trees used for associative searching. Commun. ACM 18, 509–517 (1975).

63. E. F. Pettersen et al., UCSF Chimera--a visualization system for exploratory research and analysis. J Comput Chem 25, 1605–1612 (2004).

64. C. M. Ives et al., Restoring protein glycosylation with GlycoShape. Nat Methods 21, 2117–2127 (2024).

